# IDENTIFICATION OF CANDIDATE GENES UNDERLYING A NOVEL QTL FOR ROOT LENGTH AND DIAMETER IN BARLEY IN RESPONSE TO SALINITY

**DOI:** 10.64898/2026.11.01.763390

**Authors:** Diane Mather, Apriadi Situmorang, Chris J. Brien, Megan C. Shelden

## Abstract

Salinity is a major agricultural problem that decreases growth and yield of crops and is expected to worsen with climate change. Improving salt tolerance through breeding could be facilitated by identification of genetic loci and candidate genes affecting root traits that enhance water and nutrient uptake while limiting accumulation of toxic ions.

In this study, a Clipper x Sahara 3771 doubled haploid (DH) barley mapping population was screened for root growth responses to saline conditions in a controlled environment. Quantitative trait loci (QTLs) were mapped for root traits including seminal root length, root diameter and root potassium. Major QTLs affecting seminal root length and average root diameter were detected in the distal region of the short arm of chromosome 2H, indicating that this region is a key determinant of root architecture under both control and saline conditions. The linkage map was improved by using newly designed assays for SNPs on chromosome 2H and a 2.5-Mb interval was identified for investigation of candidate genes. Bulked segregant RNA sequencing (BSR-Seq) of contrasting salt-tolerant and salt-sensitive DH lines identified differentially expressed genes associated with root growth maintenance and ion homeostasis in root tips. These findings identify specific region on chromosome 2H as a major genomic region affecting barley root architecture, especially under saline conditions, and identify differentially expressed candidate genes in that region.

## INTRODUCTION

Soil salinity is a major abiotic stress that limits growth and yield of crops worldwide. Saline soils are defined as having an electrical conductivity (EC) greater than 4 dS m^-1^ (equivalent to ∼40 mM NaCl), and can lead to a reduction in crop growth of 15 - 20 % (Munns and Tester, 2008). The severity of salinity stress on a crop depends on the timing and duration of stress, plant development stage, species and genotype. Climate change is expected to further increase soil salinisation and reduce the availability of arable land for crop production, highlighting the need to develop salt-tolerant crop varieties. Several genes encoding transporters have been identified that contribute to salt tolerance in crops, including *Nax2* in wheat (Munns *et al*., 2012), *SKC1* (*OsHKT1;5*) in rice (Kobayashi *et al*., 2017) and members of the *HAK* family in maize (Zhang *et al*., 2019). However, considerably less is known about the genetic mechanisms regulating root growth and architecture to salinity.

Barley (*Hordeum vulgare* L.) is the fourth most important cereal crop worldwide and is primarily cultivated for malting and livestock feed. Although barley is relatively salt-tolerant compared with other cereal crops, yields are constrained by salinity, as well as by drought. Salt tolerance is a complex trait, involving both osmotic and ionic components. The initial reduction in root growth following exposure to salinity occurs rapidly and is primarily associated with the osmotic component of salt stress, whereas ion toxicity develops subsequently as toxic sodium (Na^+^) and chloride (Cl^-^) ions build up in plant tissue (Munns and Tester, 2008). In both barley and wheat, seminal root growth is inhibited by NaCl in a dose-dependent manner (Rahnama *et al*., 2011; Shelden *et al*., 2013), highlighting the sensitivity of root growth to salinity and the importance of understanding the mechanisms underlying variation in root responses. Plant hormones, such as auxin, are key regulators of root growth and development and contribute to plant responses to abiotic stress (Yu *et al*., 2020).

Root system architecture is particularly important for plant adaptation to saline soils. While there are many studies examining above-ground traits for salinity tolerance, studies on root traits are limited due to the difficulties associated with measuring and phenotyping root systems. Understanding how roots respond and adapt to salinity stress is important for the development of salt-tolerant crops (Shelden and Munns, 2023). Soil salinity can affect the ability of the roots to capture water and nutrients from the soil, and results in the unwanted uptake of toxic Na^+^ and Cl^-^ ions into root cells. Maintenance of root elongation and continued acquisition of water and nutrients are important adaptive responses to salinity. In a previous study, we identified barley genotypes that differed in their ability to maintain root elongation in response to salinity (Shelden *et al*., 2013). Identifying the molecular mechanisms underlying these contrasting responses to salinity may provide new targets for breeding crops with root systems better adapted to saline soils.

Salt tolerance is a complex, multigenic trait and therefore it is difficult to identify specific gene targets for improved salt tolerance in crop species. Numerous quantitative trait loci (QTLs) associated with salinity tolerance have been reported in barley (Ellis *et al*., 2002; Rivandi *et al*., 2011; Shavrukov *et al*., 2010; Xue *et al*., 2009) but relatively few studies have identified the underlying genes for these loci. Recently, a QTL for salinity damage score and Na^+^ exclusion on chromosome 7H, was fine mapped, revealing the most likely candidate to be *HvNCX*, a sodium/calcium exchanger (Zhu *et al*., 2023). QTLs associated with root responses to salinity have been identified in several crop species including barley, maize and rice (reviewed in Karahara and Horie, 2021). In barley, salinity stress-related QTLs have been identified for root elongation (Xue *et al*., 2017), membrane potential of root epidermal cells under hypoxia (Gill *et al*., 2017), root reactive oxygen species content (Gill *et al*., 2019) and root ion content and root dry weight (Nguyen *et al*., 2013). Despite the identification of numerous salt-responsive QTLs in barley, the causal genes and molecular mechanisms underlying root growth and architecture under saline conditions remain poorly understood.

Bulked segregant analysis (BSA), involves pooling DNA from individuals with contrasting phenotypes to identify genetic markers linked to target loci controlling the trait of interest (Michelmore *et al*., 1991). Application of a bulked segregant approach for RNA sequencing (BSR-Seq) enables simultaneous identification of differentially expressed genes and sequence polymorphisms associated with the trait of interest. For example, BSR-Seq has been used to reveal candidate genes for abiotic stress tolerance in maize (*Zea mays* L.)(Du *et al*., 2017; Yan *et al*., 2023,) and sorghum (*Sorghum bicolor L.*)(Zhang *et al*., 2022). The combination of BSR-Seq with QTL analysis provides a powerful tool to understand molecular mechanisms associated with abiotic stress responses.

In this study, a Clipper x Sahara 3771 doubled haploid (DH) mapping population was used to investigate the genetic basis of root responses to salinity stress. The parental genotypes exhibit contrasting growth responses to salinity, with Clipper maintaining root elongation to a greater extent than Sahara 3771 (Shelden *et al*., 2013). Root traits were phenotyped across the DH population to identify genomic regions associated with variation in root responses to salinity. To identify candidate genes underlying major loci, BSR-Seq was performed using contrasting salt-tolerant and salt-sensitive lines. Differentially expressed genes and sequence polymorphisms identified through BSR-Seq were integrated with QTL mapping to identify candidate genes associated with salinity-responsive root traits. This combined approach provides new insights into the genetic and molecular mechanisms controlling root growth and architecture under salinity stress in barley.

## MATERIALS AND METHODS

### Plant materials, DNA marker data and genetic linkage map

Plant materials consisted of Clipper (an Australian two-rowed barley cultivar), Sahara 3771 (an Algerian six-rowed barley landrace) and 144 doubled haploid (DH) lines derived from the F_1_ generation of a cross between Clipper and Sahara 3771 (Karakousis *et al*., 2003). Seed and DNA samples of these lines, genotypic data and a “legacy” genetic linkage map of 436 markers at 261 loci (Table S1) were all provided by the Australian Centre for Plant Functional Genomics.

### Phenotypic analysis of a mapping population

Clipper, Sahara 3771 and 144 Clipper x Sahara 3771 DH lines were evaluated in a three-replicate randomized split-unit experiment. Uniform-sized seeds were selected, surface sterilised and placed on nutrient agar medium as described previously (Shelden *et al*., 2013). Seeds were germinated at 14°C in darkness before transfer to nutrient agar and grown for 72 h at 17°C in darkness. The control nutrient medium contained 0.05 mM Na^+^ supplied as NaFe-EDTA. For salt treatment, nutrient agar was supplemented with 100 mM NaCl and additional CaCl_2_ to a final activity of 1 mM (Genc *et al*., 2010). Experiments were conducted in a temperature-controlled growth cabinet (Conviron, Canada) located at The Plant Accelerator® (Australian Plant Phenomics Facility, Adelaide, Australia).

A total of 900 seedlings were evaluated across the experiment, including parental controls incorporated into each weekly batch of 50 plates, to monitor experimental consistency. Each seedling was grown individually on a 12 cm x 12 cm square petri dish, with 50 plates evaluated per week over 18 weeks.

The experiment employed a split-unit design in which each line was assigned to a pair of consecutive plates (main units) using a resolved row-and-column design. Salt and control treatments were then randomized to the two plates within each pair. As the main-unit design was resolved, each complete replicate was processed before commencement of the subsequent replicate. Weekly batches of 50 plates were handled under standardized conditions to minimize environmental variation. The main-unit design was generated using CycDesigN (VSN International, 2013), and subplot randomization was performed using the R package dae (Brien, 2017) within the R statistical computing environment (R Core Team, 2021).

#### Root and shoot phenotyping

Following 72 h of growth, seedlings were scanned using a WinRhizo® root analysis system (Regent Instruments, Canada). Total seminal root length (SRL), average root diameter (RD) and shoot length (SL) were recorded.

After scanning, shoots were excised from the roots using a razor blade and root and shoot fresh weight (RFW and SFW) were recorded. The roots were briefly rinsed in dilute 0.05 mM CaSO_4_ solution, placed in paper bags and oven-dried at 60°C for 48 h, before determination of root dry weight (RDW). Shoot dry weight (SDW) was determined using the same procedure.

Twelve phenotypic traits were measured for each line under both control and salt conditions: root fresh weight (RFW), root dry weight (RDW), total seminal root length (SRL), average root diameter (RD), shoot length (SL), shoot fresh weight (SFW), shoot dry weight (SDW), root and shoot Na^+^ and K^+^ concentration (*Section 2.4*).

#### Determination of Na^+^ and K^+^ content

Root and shoot Na^+^ and K^+^ concentrations were determined as previously described (Munns *et al*., 2010; Shelden *et al*., 2013). Dried tissue samples were digested in 1% (v/v) nitric acid at 85°C for 2 h and subsequently diluted with Milli-Q water as required.

Na^+^ and K^+^ concentrations were quantified by atomic absorption spectrometry (AA-7000, Shimadzu Corporation, Japan) in flame mode as described previously (Shelden *et al*., 2020). Standard curves were prepared using 1000 ppm Na and K standard solutions (SIGMA, Australia). Caesium chloride was added to samples and standards to a final concentration of 0.1% (w/v) to minimise ionisation interference during analysis.

### Statistical analysis of phenotypic data

Phenotypic means were adjusted for the spatial variation in the growth room, using a linear-mixed-model analysis performed for each trait using the R package ASReml (Butler *et al*., 2009) and ASRremlPlus (Brien, 2018) within the R statistical environment (R Core Team, 2021). This model included terms for two-factor factorial model of the effects of genotype (lines), treatment (conditions), and for large-scale spatial variation between replicates, weeks within replicates and MainUnit within Weeks. It also allowed for the variances of the roots from control plates to differ from those from salt plates.

For each trait, REML ratio tests (*_α_* = 0.05) were used to determine whether the variances differed significantly between the treatments, and models were modified accordingly. Model assumptions were assessed by residual-versus-fitted values plots, and normal probability plots of the residuals.

Wald F-tests were conducted to assess significance of genotype (lines) x treatment (conditions) interactions. Where interactions were not significant, the main effects of genotype and treatment were evaluated. Best linear unbiased estimates (BLUEs) were obtained for each genotype x treatment combination and used for QTL analysis, selection of BSR-Seq bulks, and subsequent phenotypic analyses.

### Selection of bulks for BSR-Seq analysis

Twenty DH lines were selected from the mapping population for bulked segregant RNA sequencing (BSR-Seq). Ten of these lines were classified as sensitive to salinity because their roots were significantly shorter in the salt treatment compared to the control treatment. The other ten lines were classified as tolerant to salinity because their roots were not significantly shorter in the salt treatment compared to the control treatment. Each of the sensitive lines was allocated to one of two bulks (S1 and S2) and each of the tolerant lines was allocated to one of two other bulks (T1 and T2) (Table S2).

Two plants of each selected line were grown in duplicates under control and salt conditions, in each of three independent experiments, using the methods described in Section 2.2. After 72 h of treatment, 5 mm root tips were excised from the three longest seminal roots of each seedling using a razor blade under a dissecting microscope. Root tips from the two seedlings of each DH line were pooled to form one sample to obtain enough tissue for total RNA extraction. Equal amounts of tissue from the 4-6 DH lines comprising each bulk were then combined to generate a single bulked biological replicate. Samples were immediately transferred to pre-chilled microcentrifuge tubes, snap frozen in liquid N, and stored at −80°C until RNA extraction. Four bulks were analysed under two treatments (control and salt; total eight bulks) with three biological replicates per treatment, giving a total of 24 RNA samples.

### RNA extraction, library construction, and sequencing

Total RNA was extracted using the Spectrum™ Plant Total RNA Kit (Sigma-Aldrich, Missouri, USA) according to the manufacturer’s instructions. Residual genomic DNA was removed using a Turbo DNase™ kit (Invitrogen™, MA, USA). RNA quantity and quality were assessed using an Agilent 2100 Bioanalyzer (SA Pathology, Adelaide, SA, Australia). All samples had an RNA integrity (RIN) > 7.4.

RNA sequencing was performed at the Australian Genomics Research Facility (AGRF, Melbourne, VIC). Twenty-four cDNA libraries were prepared and sequenced on an Illumina NovaSeq 6000 platform using NovaSeq 6000 v1.5 chemistry to generate 2 x 100 bp paired-end reads.

### Differential gene expression analyses

RNA-seq data pre-processing was carried out using an in-house analysis workflow, using MultiQC for quality reporting (Ewels *et al*., 2016), alignment to the barley reference genome assembly using STAR (Dobin *et al*., 2013), and gene expression quantification using featureCounts (Liao *et al*., 2014). Alignment rates exceeded 90% for all libraries and sequencing depth exceeded 45 million paired-end reads per library. Differential gene analysis was performed in R (R Core Team, 2021) using the edgeR (Robinson *et al*., 2009) and limma-voom (Law *et al*., 2014; Ritchie *et al*., 2015) packages. Genes with counts per million (CPM) > 1 in at least three libraries were retained for analysis resulting in 20,604 expressed genes. Library sizes were normalised using the trimmed mean of M-values (TMM) method implemented in edgeR. Differential gene expression was performed using limma-voom and assessed using empirical Bayes moderated t-tests implemented in limma. Genes with a false discovery rate (FDR)-adjusted *p*-value < 0.05 and log_2_fold-change |log_2_FC)| > 1 were considered differentially expressed.

Gene Ontology (GO) and Kyoto Encyclopedia of Genes and Genomes (KEGG) pathway enrichment analyses of salt-responsive DEGs were performed using ShinyGO V0.86.0 (Ge *et al*., 2020), with *Hordeum vulgare* specified as the reference species. GO and KEGG pathways with an FDR <0.05 were considered significantly enriched.

GO enrichment analysis of genes exhibiting a root-length-dependent response to salinity was performed separately using g:Profiler, with *Hordeum vulgare* specified as the reference organism (Kolberg *et al*., 2023). Enrichment was assessed for the molecular function (GO:MF), biological process (GO:BP), and cellular component (GO:CC) categories. GO terms with adjusted P < 0.05 were considered significantly enriched.

### Variant calling and annotation using BSR-Seq data

Single nucleotide polymorphisms (SNPs) and short insertion/deletion variants (indels) were identified from RNA-seq reads generated from the four contrasting bulks. Reads had been previously aligned to the barley Morex V3 reference genome during transcriptome analysis. Variant discovery was performed according to the Genome Analysis Toolkit (GATK) best practice workflow for RNA-seq short variant discovery. Briefly, aligned reads were processed using SplitNCigarReads prior to variant calling. Variants were called for each library using GATK HaplotypeCaller in gVCF mode, followed by joint genotyping using CombineGVCFs and GenotypeGVCFs.

Variants were filtered according to GATK recommendations for RNA-seq data. Variants were annotated using SnpEff (Cingolani et al., 2012) with the *Hordeum vulgare* L. database compatible with the Morex V3 genome assembly. Only annotated variants that passed quality and coverage filtering criteria were retained for downstream analyses.

### Quantitative trait loci analysis and SNP genotyping

Quantitative trait locus analysis was conducted using simple interval mapping as implemented in the R package R/qtl (Broman et al. 2009), with BLUEs from the phenotypic analysis used as trait values for the DH lines. Logarithm of odds (LOD) test statistic values were compared to threshold values determined with 10,000 permutations to achieve a genome-wide significance level of 0.05.

After QTL analysis revealed that both root length and root diameter were affected by the terminal region of the short arm of chromosome 2H, genotype results for variants on that chromosome arm were reviewed to identify single-nucleotide polymorphisms (SNPs) for which assays could be designed to improve the genetic map. Competitive allele-specific PCR (KASP) assays (He et al. 2014) were designed for 16 SNPs. The positions of these SNPs in Version 3 of the Morex barley genome assembly (Mascher et al. 2021) and the sequences of the KASP primers are provided in Table S3.

The KASP assays were applied to Clipper, Sahara 3771 and 144 DH lines using an automated SNPline system (LGC Biosearch Technologies, Teddington, UK) according to the manufacturer’s instructions. Genotypic data for the 16 SNPs were combined with genotypic data for legacy markers that had previously been mapped on chromosome 2H. Where SNPs co-segregated with legacy markers, the SNPs were retained and the legacy markers were removed. Based on the combined dataset, marker loci on chromosome 2H were ordered and genetic distances calculated using the MSTmap algorithm (Wu et al. 2008) as implemented in the R/ASMap package (Taylor et al. 2017). QTL analysis was repeated using the SNP-enriched linkage map, with significance thresholds established based on 10,000 permutations for root length and root diameter, each under both control and salt conditions.

### Filtering of BSR-Seq variants in a candidate region on chromosome 2H

Variants within a candidate region (500,000 to 3,000,000 bp on chromosome 2H) were reviewed to identify variants as candidate polymorphisms for QTL effects. For variants with high or moderate SNPeff impact ratings, genotype results for the treatment combination (two sets of T lines and two sets of S lines, each grown in control and salt treatments) were considered. Each treatment combination was classified as fixed for a particular allele if all three of its replicate samples had been called as fixed for that allele, or if two replicate samples had been called as fixed for that allele and the remaining sample not called (null result) or called as mixed (heterozygous). Variants were then excluded from further consideration if (1) none of the T treatment combinations were called as fixed for any allele; (2) none of the S treatment combinations were called as fixed for any allele; (3) more than one fixed genotype was called among the four T treatment combinations or among the four S treatment combinations; or (4) the same allele was called as fixed in both S and T treatment combinations. This filtered the variants to include only those for which at least one of the four T treatment combinations seems to be fixed for one allele (deduced to come from Clipper) and at least one of the four S treatment combinations seems to be fixed for a different allele (deduced to come from Sahara 3771), with no strong inconsistency among replicate samples, among the T treatment combinations or among the S combinations.

## RESULTS

### Phenotyping of DH mapping population for salt tolerance

Phenotypic variation among the DH lines and parental genotypes under control and salt conditions is summarised in Table S4. Frequency distributions for seminal root length, average root diameter and root Na^+^ content were approximately normally distributed (Figure 1). The remaining traits also exhibited approximately normal distributions (Figure S1). For most traits, the population mean was intermediate between the parental values. However, for average root diameter (ARD), root Na^+^ content (RNaC) and shoot length (SL), the population mean fell outside the parental range.

**Figure 1.**
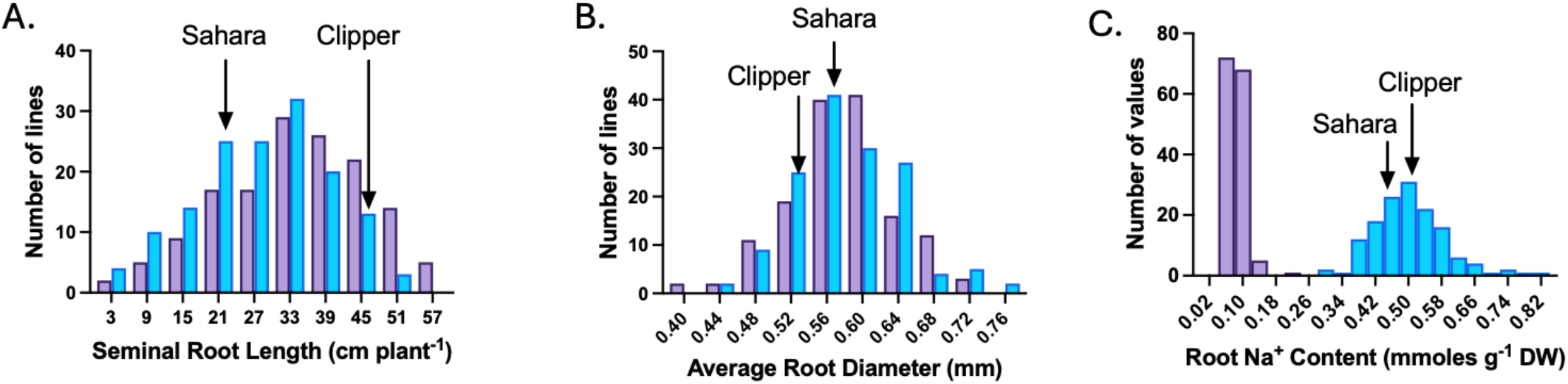
Root growth under control and salt conditions (Clipper, Sahara 3771, tolerant, sensitive DH line). Frequency distributions of (A) seminal root length (SRL), (B) average root diameter (ARD) and (C) root Na^+^ content for the Clipper x Sahara 3771 DH population under control (purple) and salt treatment (blue). Arrows indicate the parental values under salt-treatment. Phenotypic values represent BLUEs calculated from three independent experiments.

### Phenotypic validation of selected salt-tolerant and salt-sensitive lines

Twenty lines were classified as salt-tolerant (T1 and T2) and salt-sensitive (S1 and S2) and selected for further analysis based on their root phenotype under saline conditions in the phenotyping experiment described in Section 3.1. The root phenotypic responses of the selected lines are shown in Figure 2. Both T1 and T2 maintained seminal root length in response to salt treatment relative to control at 88 and 91%, respectively, with no significant difference between salt-treated and control plants or between T1 and T2. Seminal root length was significantly higher for T1 and T2 compared to S1 and S2 for both control and salt treatment (Figure 2A). S1 and S2 relative root length (salt relative to control) was 71 and 73%, respectively. S1 and S2 salt-treated roots were significantly shorter than their corresponding controls.

**Figure 2.**
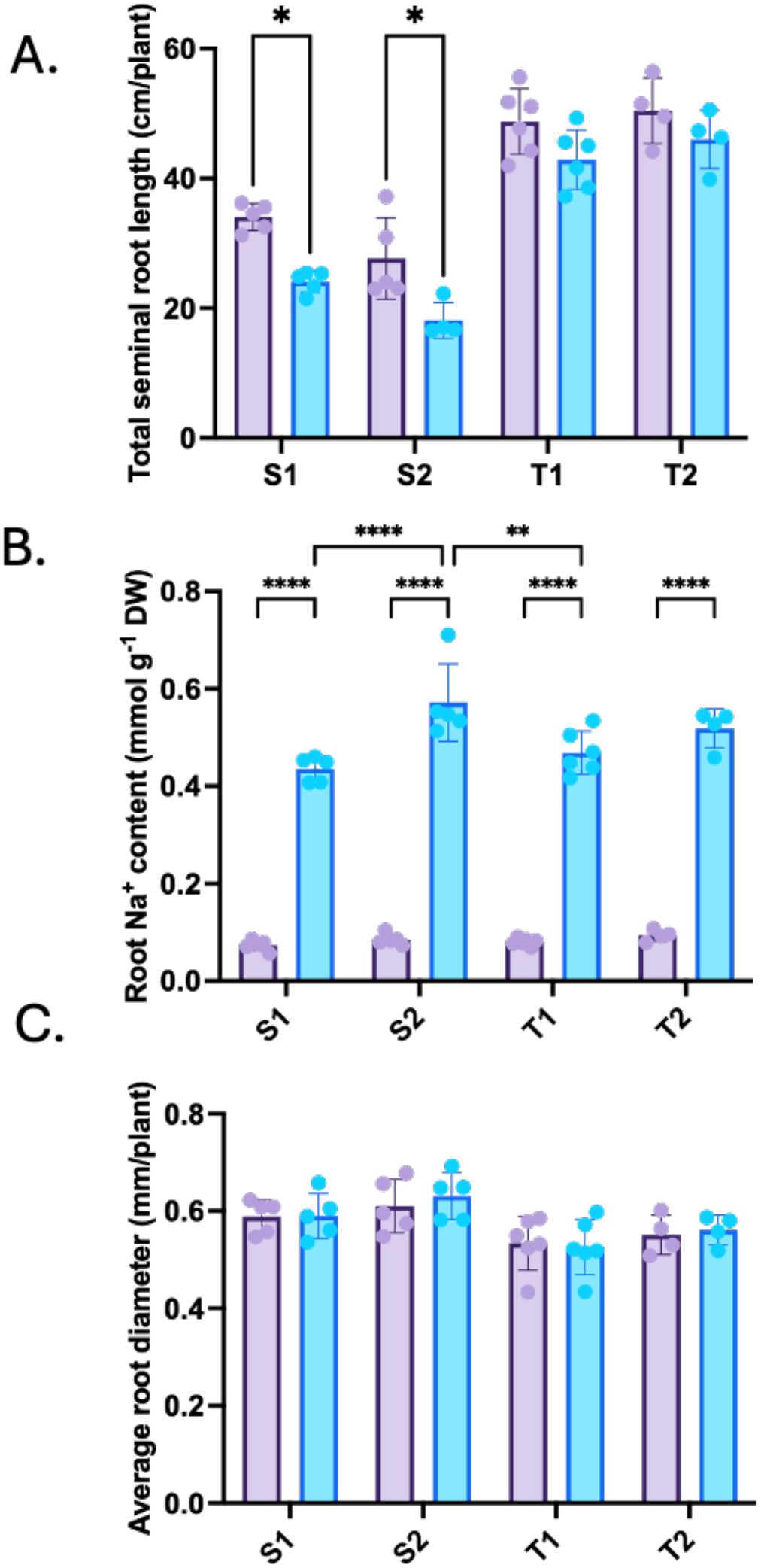
Graphs showing the distribution of root traits in salt-sensitive (S1 and S2) and salt-tolerant (T1 and T2) barley lines used as bulks for BSR-seq. Root traits were measured after 3 d of growth on control (purple) or 100 mM NaCl (blue) nutrient media. (A) Total seminal root length, (B) root Na^+^ content and (C) average root diameter. Each bar represents the mean of the BLUE values for each bulk, n = 5 (S1), 5 (S2), 6 (T1), 4 (T2) ± SEM. Asterisks indicate significant differences between groups (* p < 0.05, ** p < 0.01, **** p < 0.0001; two-way ANOVA, Tukey’s multiple comparison test).

For all groups, salt-treated roots had significantly higher root Na^+^ content (5.5 to 7.1-fold) compared to their respective controls (Figure 2B). No significant difference in root Na^+^ content was observed between T1 and T2, however S1 and S2 were significantly different to each other (Figure 2C). The average root diameter was lower in T1 and T2 compared to S1 and S2 (Figure 2C). These selected lines were then used for bulk segregant RNA sequencing.

### BSR-Seq analysis of tolerant and sensitive lines

RNA sequencing generated a total of 733.5 M reads across the 24 libraries representing the T1, T2, S1 and S2 pools under control and salt conditions, with three biological replicates per treatment combination. Genes were considered differentially expressed (DE) if their FDR-adjusted *p*-value was <0.05 and |log_2_FC| > 1. Principal component analysis (PCA) of the RNA-seq data showed “salt treatment” separated the samples across PC2, indicating that salt treatment is one of the main sources of variation in the data, accounting for approximately 10% of the variation in gene expression between samples (Figure 3A). PC1 explained 28% of the transcriptional variation and separated tolerant and sensitive bulks.

**Figure 3.**
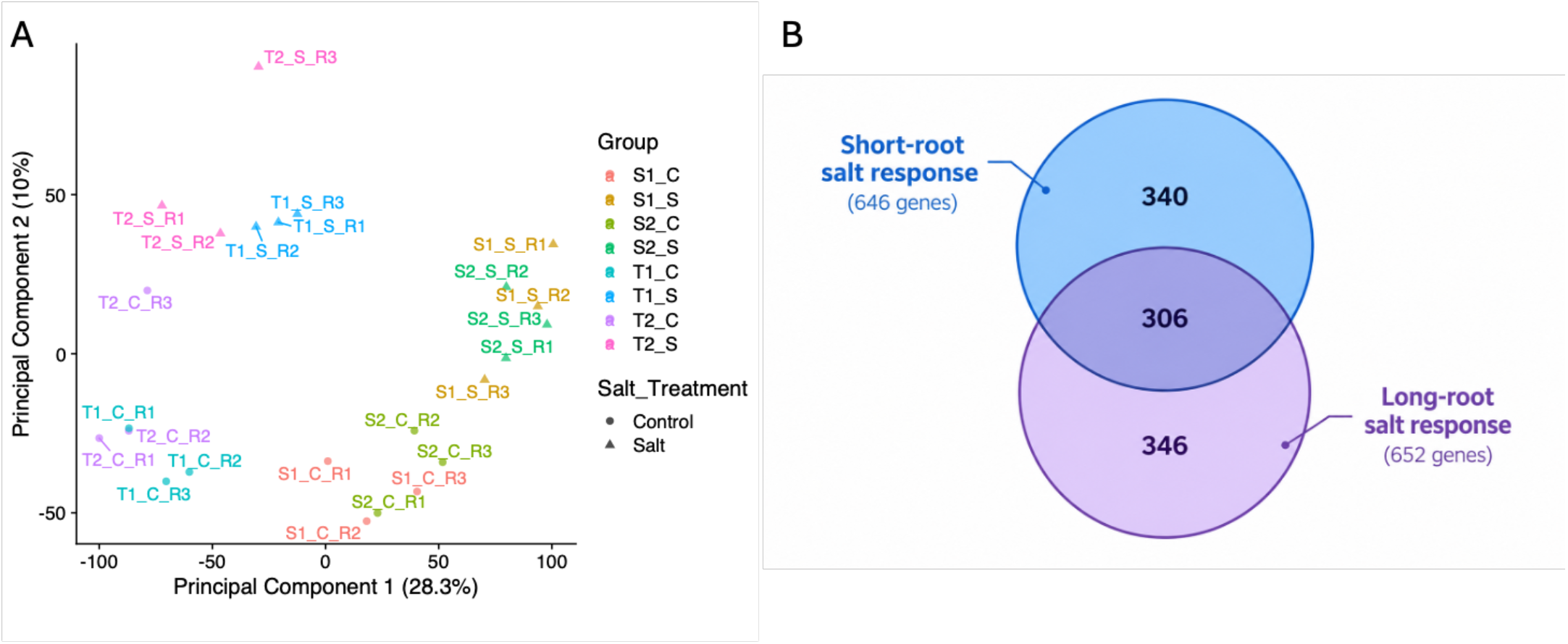
Differential expression in barley root tips. (A) Principal component analysis (PCA) plot of RNA-seq TMM read counts (gene expression) showed separation of control (C) and salt-treated (S) samples on PC1 and separation of tolerant (T1 and T2) and sensitive (S1 and S2). Each point represents one bulked sample with R1, R2 and R3 representing samples from three independent experiments. (B) Venn diagram showing the number of differentially expressed genes (DEGs) in long roots (T1 and T2) and short roots (S1 and S2) in response to salt stress. The number in overlapping areas represent the number of shared DEGs.

#### Salt-responsive genes in long and short roots

In response to salinity stress, 652 and 646 genes were differentially expressed in the tolerant (long-root) and sensitive (short-root) bulks, respectively. In the tolerant bulks, 606 genes were upregulated and 46 were downregulated, whereas the sensitive bulks exhibited 569 upregulated and 77 downregulated genes, in response to salt treatment. The tolerant and sensitive bulks shared 306 salt-responsive genes, while each bulk also exhibited distinct transcriptional responses associated with their root-length phenotype (Figure 3B).

KEGG enrichment analysis of salt-induced genes using ShinyGO identified several pathways enriched in both tolerant and sensitive bulks, including glutathione metabolism, plant MAPK signalling, phenylpropanoid biosynthesis and biosynthesis of secondary metabolites. In the tolerant bulks, glutathione metabolism showed the highest fold enrichment, followed by plant MAPK signalling and phenylpropanoid biosynthesis (Figure 4A). Plant hormone signal transduction was also significantly enriched. In contrast, the sensitive bulks showed strong enrichment of alpha linolenic metabolism, and zeatin biosynthesis (Figure 4B). Genes differentially expressed between tolerant and sensitive bulks were enriched for pathways associated with flavone and flavanol biosynthesis, carotenoid biosynthesis, flavonoid biosynthesis, nitrogen metabolism and phenylpropanoid synthesis (Figure 4C).

**Figure 4.**
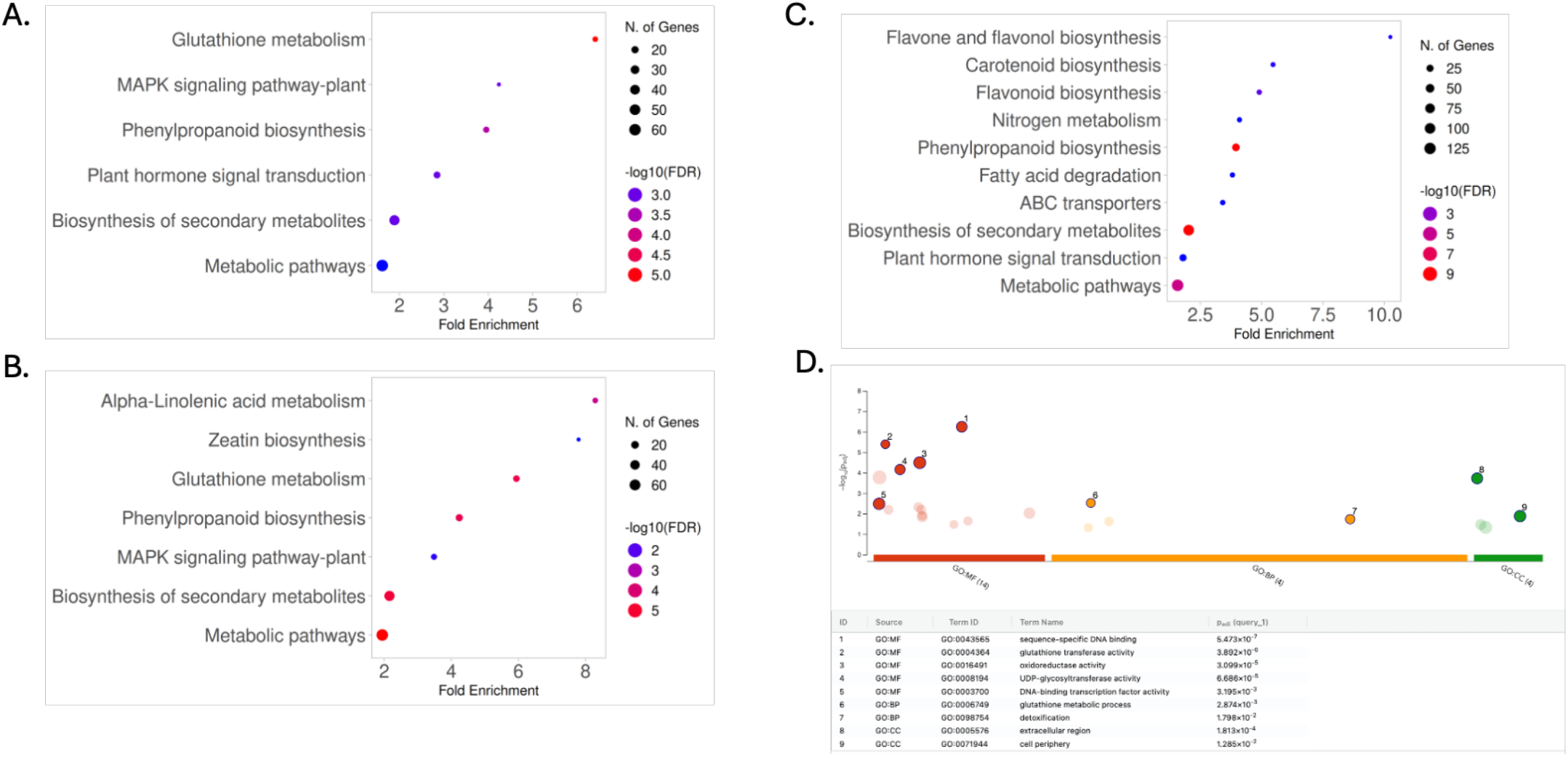
Profiling of DEGs in barley roots under control and salt stress in the tolerant and sensitive bulks. KEGG enrichment analysis of DEGs using ShinyGO tool. The enriched KEGG pathway in (A) tolerant and (B) sensitive bulks under salt stress. (C) DEGs enriched in tolerant bulks compared to sensitive bulks. False discovery rate (FDR; coloured dot) represents the degree of enrichment. The size of the dot indicates the number of DEGs involved in the pathway. (D) GO analyses of DEGs in root length dependent salt response.

#### Root-length dependent salt-response

A total of 83 differentially expressed genes (DEGs) exhibited a root-length-dependent response to salt stress, indicating that the expression of these genes was influenced by both factors and their combined effects. Of these, 66 DEGs showed positive interaction effects and 17 showed negative interaction effects (Table S5). GO enrichment analysis identified sequence-specific DNA binding (GO:0043565; adjusted *P* = 6.22 × 10⁻¹⁰) and DNA-binding transcription factor activity (GO:0003700; adjusted *P* = 1.00 × 10⁻⁷) as the most significantly enriched terms. Regulation of RNA biosynthetic process (GO:2001141; adjusted *P* = 1.18 × 10⁻⁴) and systemic acquired resistance (GO:0009627; adjusted *P* = 2.03 × 10⁻²) were also significantly enriched (Figure 4D). Overall, genes exhibiting a root-length-dependent response to salinity were predominantly enriched for DNA-binding and transcriptional regulatory functions.

### Quantitative Trait Loci

With QTL analysis using the legacy linkage map, QTLs were mapped for root morphology traits, biomass traits, and ion homeostasis traits under control and/or saline conditions (Tables S6 C S7). Significant LOD scores ranged from 2.5 to 10.5.

#### Root morphology

The results of the QTL analysis for root traits indicate that the terminal region of the short arm of chromosome 2H is important in determining root morphology in both control and saline conditions (Table S6), with alleles from Sahara 3771 favouring short, thick roots and Clipper alleles favouring long, thin roots.

#### Biomass

Near *Vrs1,* the major gene on chromosome 2H that determines whether barley spikes have two or six rows of spikelets (Saisho *et al*., 2009), QTLs were detected for root and shoot dry weight. Another QTL for shoot dry weight was detected on chromosome 1H, near the *HvNax4* sodium exclusion locus (Rivandi *et al*., 2011). In all cases, the allele from Sahara 3771 was associated with lower dry weight.

#### Ion homeostasis

QTLs associated with Na^+^ and/or K^+^ content in roots and shoots were detected on chromosomes 1H, 2H and 6H (Table S7). On chromosome 1H, there were significant LOD scores for Na^+^ content and K^+^/Na^+^ ratio, but the QTL peaks for those traits were not very distinct. On chromosome 2H, there are QTLs at which the Sahara 3771 allele increases shoot K^+^ concentration (at 144 cM) or root Na^+^ concentration (at 151 and 198 cM). On chromosome 6H, there is a minor QTL at which the Sahara 3771 allele reduces root Na^+^ concentration in saline conditions.

#### Refining the linkage and QTL maps for chromosome 2H

To further investigate the root morphology QTLs on chromosome 2H, newly designed KASP assays for 16 SNPs (all discovered with BSR-Seq) were used to enrich the linkage map. All 16 assays discriminated between the two parental lines and detected segregation in the mapping population. All 16 SNPs were genetically mapped on the short arm of chromosome 2H (Table S8).

With QTL analysis using the SNP-enriched map, QTLs for seminal root length and average root diameter were mapped on the short arm of chromosome 2H (Table 1, Figure 5). Under saline conditions, both traits had very distinct QTL peaks at 2.1 cM (Figure 5B). Under control conditions, the QTL peaks were less distinct and had lower LOD values. Based on these results, 2.1 cM was taken to be the best estimate for the position of a QTL that affects both seminal root length and average root diameter, particularly under saline conditions. Based on the physical positions of four SNPs that mapped at 2.1 cM (Figure 6) and considering the sharp drops in LOD values immediately distal and proximal to the QTL peaks (Figure 5B), the 2.5-Mb region of the 2H barley pseudomolecule between 500,000 bp and 3,000,000 bp provides an approximate 95% confidence interval for the physical position of the causal gene(s). Consistent with the detection of a QTL at 2.1 cM, all 10 DH lines in the two tolerant groups carry Clipper alleles at the 2.1-cM and 2.8-cM positions and all 10 lines in the two sensitive groups carry Sahara 3771 alleles at those positions (Table S9).

**Figure 5.**
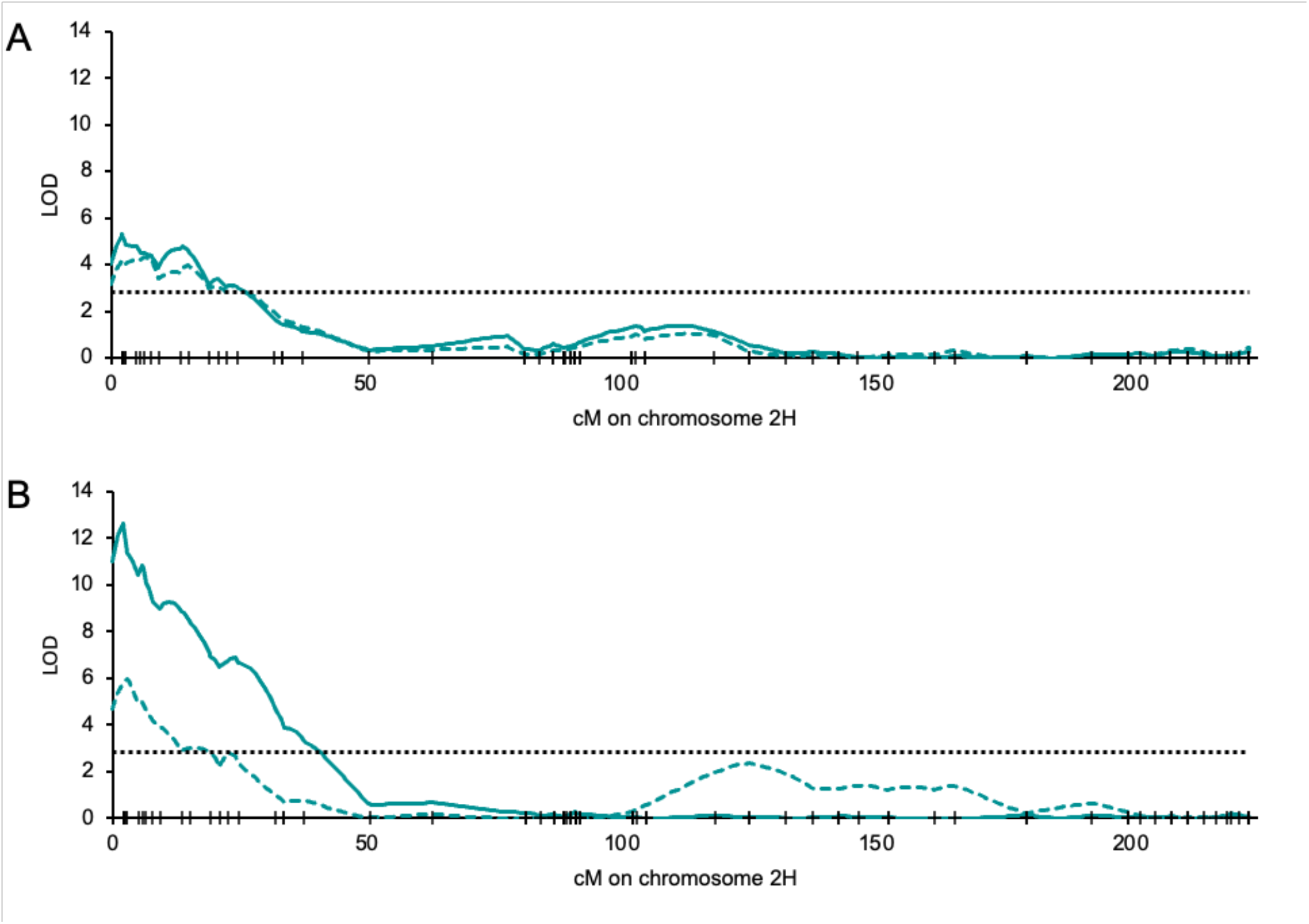
LOD test statistic scans for barley chromosome 2H for (A) total seminal root length and (B) average root diameter of Clipper x Sahara 3771 doubled haploid lines grown for 72 hours on control (dashed line) or saline (solid line) nutrient medium. Tick marks on the horizontal axis show the genetic positions at which markers were mapped. The horizontal dashed line shows the genome-wide significance threshold (LOD= 2.8).

**Figure 6.**
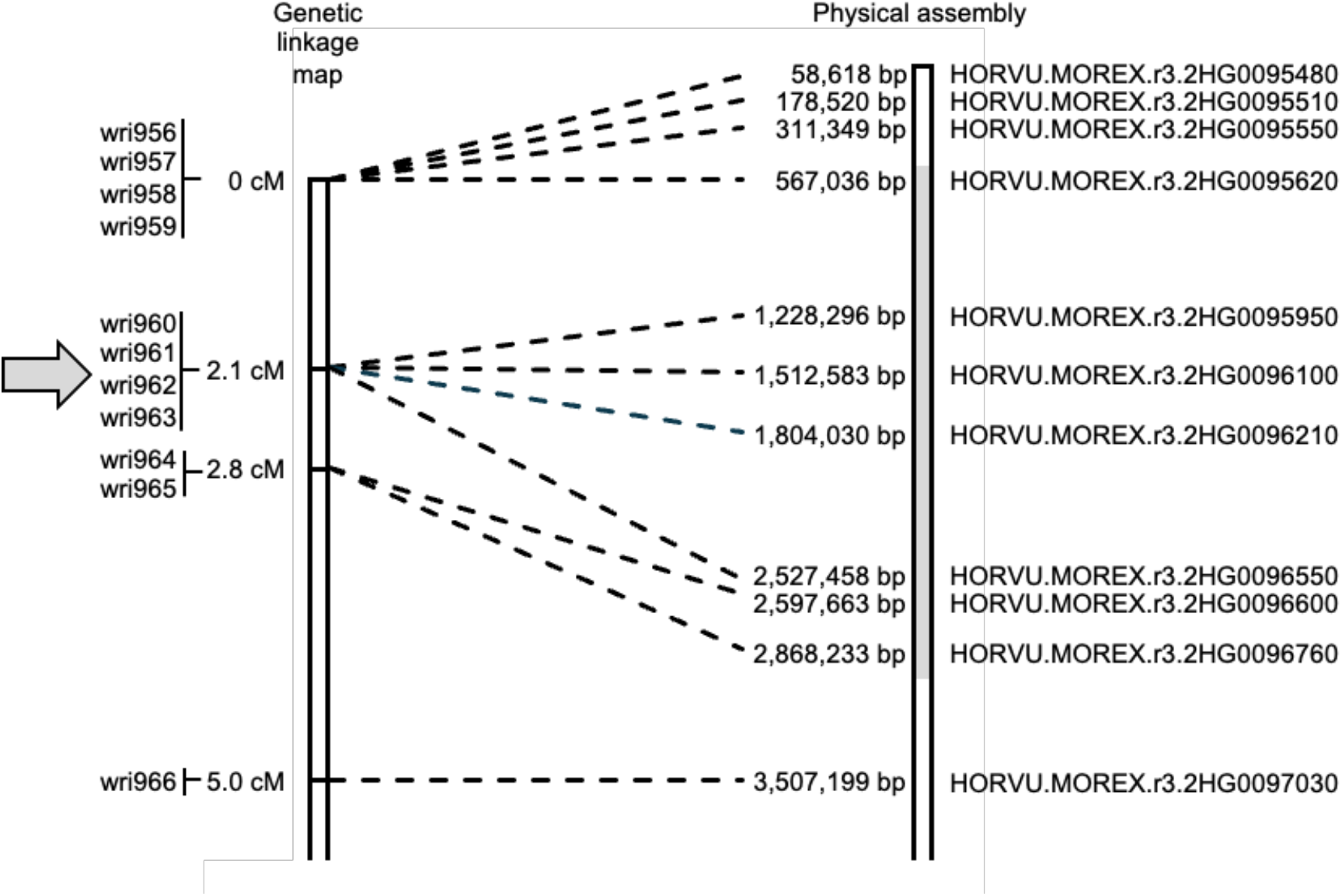
Genetic linkage map (on the left) showing the positions in the distal 5 cM of the short arm of chromosome 2HS at which SNPs were mapped using assays wri956 through wri966 and (on the right) the physical positions of those SNPs on the 2H pseudomolecule of Version 3 of the Morex barley genome assembly, and the names of the predicted genes in which the SNPs are located. The estimated genetic position of the QTL affecting total seminal root length and average root diameter under saline conditions is marked with an arrow and the candidate region of the pseudomolecule is shaded.

**Table 1.**
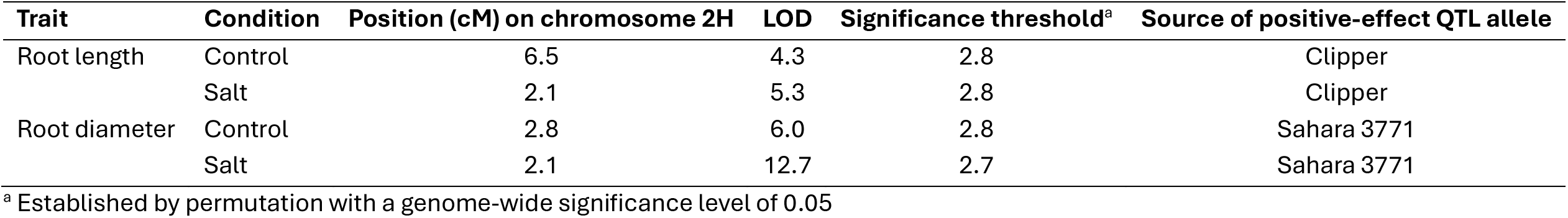
Quantitative trait loci detected for length and diameter in a Clipper/Sahara 3771 barley population, using a linkage map that includes the positions of 16 SNP markers on the short arm of chromosome 2H

### Candidate genes for a QTL

Within the 2.5 Mb candidate region on the short arm of chromosome 2H, there are 76 high-confidence gene models on the Morex reference genome (Table S10). Ten of these genes were differentially expressed between the tolerant lines (all of which carried Clipper alleles at 2.1 and 2.8 cM) and the sensitive lines (all of which carried Sahara 3771 alleles at 2.1 and 2.8 cM). Two of these genes and two other genes were differentially expressed in response to salt stress in either the tolerant or sensitive lines (Table 2). Within the same candidate region, 3,876 sequence variants were identified, of which 177 variants in 31 genes (Table 3, Table S11) were identified as high- or moderate-impact variants that are likely to be polymorphic between Clipper and Sahara 3771.

**Table 2.** High-confidence predicted genes in the candidate region between 500,000 – 3,000,000 bp on chromosome 2H that are differentially expressed.

|  |  |  |  | Log<sub>2</sub> FC<sup>a</sup> |  |  |
|---|---|---|---|---|---|---|
| Start (bp) | End (bp) | Gene ID | Description | Tolerant vs Sensitive | Sensitive S vs C | Tolerant S vs C |
| 1,227,414 | 1,231,905 | HORVU.MOREX.r3.2HG0095950 | NBS-LRR disease resistance protein-like protein | 1.14 |  |  |
| 1,512,496 | 1,513,768 | HORVU.MOREX.r3.2HG0096100 | 3'-N-debenzoyl-2'-deoxytaxol N-benzoyltransferase | -2.85 |  |  |
| 1,524,394 | 1,525,030 | HORVU.MOREX.r3.2HG0096120 | WEB family protein (DUF827) |  | 1.21 |  |
| 1,818,365 | 1,818,761 | HORVU.MOREX.r3.2HG0096230 | Phosphatidylinositol 3- and 4-kinase family protein |  |  | 1.00 |
| 2,503,905 | 2,506,708 | HORVU.MOREX.r3.2HG0096520 | Pentatricopeptide repeat-containing protein (PPR) | -3.35 |  | -2.38 |
| 2,516,439 | 2,517,917 | HORVU.MOREX.r3.2HG0096540 | Glycosyltransferase | -1.89 |  |  |
| 2,523,096 | 2,530,883 | HORVU.MOREX.r3.2HG0096550 | ABC transporter B family-like protein | -4.33 |  |  |
| 2,697,963 | 2,703,943 | HORVU.MOREX.r3.2HG0096670 | Mediator of RNA polymerase II transcription subunit 15a | 3.18 |  |  |
| 2,770,503 | 2,772,104 | HORVU.MOREX.r3.2HG0096710 | Cytochrome P450 | -2.84 | 1.39 |  |
| 2,813,143 | 2,816,820 | HORVU.MOREX.r3.2HG0096730 | Agenet domain, putative | 1.32 |  |  |
| 2,872,655 | 2,876,682 | HORVU.MOREX.r3.2HG0096770 | cellulose synthase, putative (DUF1644) | 3.18 |  |  |
| 2,893,692 | 2,895,546 | HORVU.MOREX.r3.2HG0096800 | Flavin-containing monooxygenase (YUCCA 5) | -1.28 |  |  |

<sup>a</sup> Log2FC, Tolerant – T1 and T2 bulk, Sensitive – S1 and S2 bulk, S – salt treatment, C – control.

**Table 3.** High-confidence predicted genes in the candidate region between 500,000 – 3,000,000 bp on chromosome 2H for which transcriptomic sequence data indicate that Clipper and Sahara 3771 alleles differ for variants of types expected to have high or moderate impact.

| Gene ID | Description | High-impact variants | Moderate-impact variants |
|---|---|---|---|
| HORVU.MOREX.r3.2HG0095600 | NBS-LRR resistance-like protein |  | 1 missense |
| HORVU.MOREX.r3.2HG0095610 | YIF1b |  | 1 missense |
| HORVU.MOREX.r3.2HG0095620 | Curved DNA binding protein |  | 2 in-frame indel |
| HORVU.MOREX.r3.2HG0095670 | Cysteine synthase |  | 1 missense |
| HORVU.MOREX.r3.2HG0095710 | Disease resistance protein (NBS-LRR class) family |  | 2 missense |
| HORVU.MOREX.r3.2HG0095930 | NBS-LRR disease resistance protein-like protein |  | 1 missense |
| HORVU.MOREX.r3.2HG0095950 | NBS-LRR disease resistance protein-like protein | 2 stop codon gain/loss; 2 frameshift | 38 missense |
| HORVU.MOREX.r3.2HG0095960 | NBS-LRR disease resistance protein-like protein | 3 frameshift | 15 missense |
| HORVU.MOREX.r3.2HG0095970 | NBS-LRR disease resistance protein-like protein | 1 stop codon gain/loss; 1 frameshift | 9 missense; 1 in-frame indel |
| HORVU.MOREX.r3.2HG0096080 | NBS-LRR disease resistance protein-like protein | 3 stop codon gain/loss; 2 frameshift | 1 in-frame indel |
| HORVU.MOREX.r3.2HG0096100 | 3'-N-debenzoyl-2'-deoxytaxol N-benzoyltransferase |  | 8 missense |
| HORVU.MOREX.r3.2HG0096180 | Histone-lysine N-methyltransferase | 1 frameshift |  |
| HORVU.MOREX.r3.2HG0096210 | Phosphotransferase | 1 frameshift |  |
| HORVU.MOREX.r3.2HG0096220 | WEB family protein |  | 1 missense |
| HORVU.MOREX.r3.2HG0096230 | Phosphatidylinositol 3- and 4-kinase family protein |  | 1 missense |
| HORVU.MOREX.r3.2HG0096290 | Late embryogenesis abundant (LEA) hydroxyproline-rich glycoprotein family member |  | 1 missense |
| HORVU.MOREX.r3.2HG0096350 | 3'-N-debenzoyl-2'-deoxytaxol N-benzoyltransferase | 1 frameshift | 1 missense |
| HORVU.MOREX.r3.2HG0096400 | Cortactin-binding protein 2 | 1 stop codon gain/loss | 6 missense;1 in-frame indel |
| HORVU.MOREX.r3.2HG0096440 | tRNA-dihydrouridine(47) synthase [NAD(P)(+)] |  | 1 in-frame indel |
| HORVU.MOREX.r3.2HG0096450 | Peptidyl-prolyl cis-trans isomerase |  | 1 missense |
| HORVU.MOREX.r3.2HG0096460 | NBS-LRR-like resistance protein |  | 1 missense |
| HORVU.MOREX.r3.2HG0096520 | Pentatricopeptide repeat-containing protein |  | 1 missense |
| HORVU.MOREX.r3.2HG0096530 | Arginine/serine-rich splicing factor (putative) |  | 1 missense |
| HORVU.MOREX.r3.2HG0096540 | Glycosyltransferase |  | 4 missense |
| HORVU.MOREX.r3.2HG0096550 | ABC transporter B family-like protein |  | 4 missense |
| HORVU.MOREX.r3.2HG0096600 | Pyruvate dehydrogenase E1 component subunit alpha |  | 1 in-frame indel |
| HORVU.MOREX.r3.2HG0096610 | StAR-related lipid transfer protein |  | 1 missense |
| HORVU.MOREX.r3.2HG0096760 | TOM1-like protein 2 |  | 3 missense |
| HORVU.MOREX.r3.2HG0096780 | Evolutionarily conserved C-terminal region 2 |  | 9 missense |
| HORVU.MOREX.r3.2HG0096800 | Flavin-containing monooxygenase |  | 2 missense |
| HORVU.MOREX.r3.2HG0096810 | Werner-Syndrome-like exonuclease |  | 1 missense |

Based on genomic location, differential expression between contrasting bulks, transcriptional responses to salinity, sequence polymorphisms and functional annotation, six genes were prioritised as candidate genes (Table 4). HORVU.MOREX.r3.2HG0096800 is predicted to encode a flavin-containing monooxygenase homologous to YUCCA5, a key enzyme in auxin biosynthesis, and HORVU.MOREX.r3.2HG0096550 is predicted to encode an ABCB transporter protein. Both genes are downregulated in lines carrying the Clipper allele compared to the Sahara allele (flavin-containing monooxygenase log_2_FC = −1.27; ABCB transporter log_2_FC = −4.33; Figure 7, Table 2) and both have missense polymorphisms between Clipper and Sahara 3771 (Table 3). Four additional candidates, with possible functions in auxin biosynthesis, hormone transport, cell-wall biosynthesis or stress signalling are HORVU.MOREX.r3.2HG0096770, HORVU.MOREX.r3.2HG0096540, HORVU.MOREX.r3.2HG0096710 and HORVU.MOREX.r3.2HG0096520, which are predicted to encode a cellulose synthase, a glycosyltransferase, a cytochrome P450 enzyme and a pentatricopeptide repeat-containing protein (PPR), respectively. Like the auxin-associated candidates, these genes were differentially expressed between contrasting bulks (Table 2). Three of them (all except the cellulose synthase) also have missense polymorphisms between the parental alleles (Table 3).

**Figure 7.**
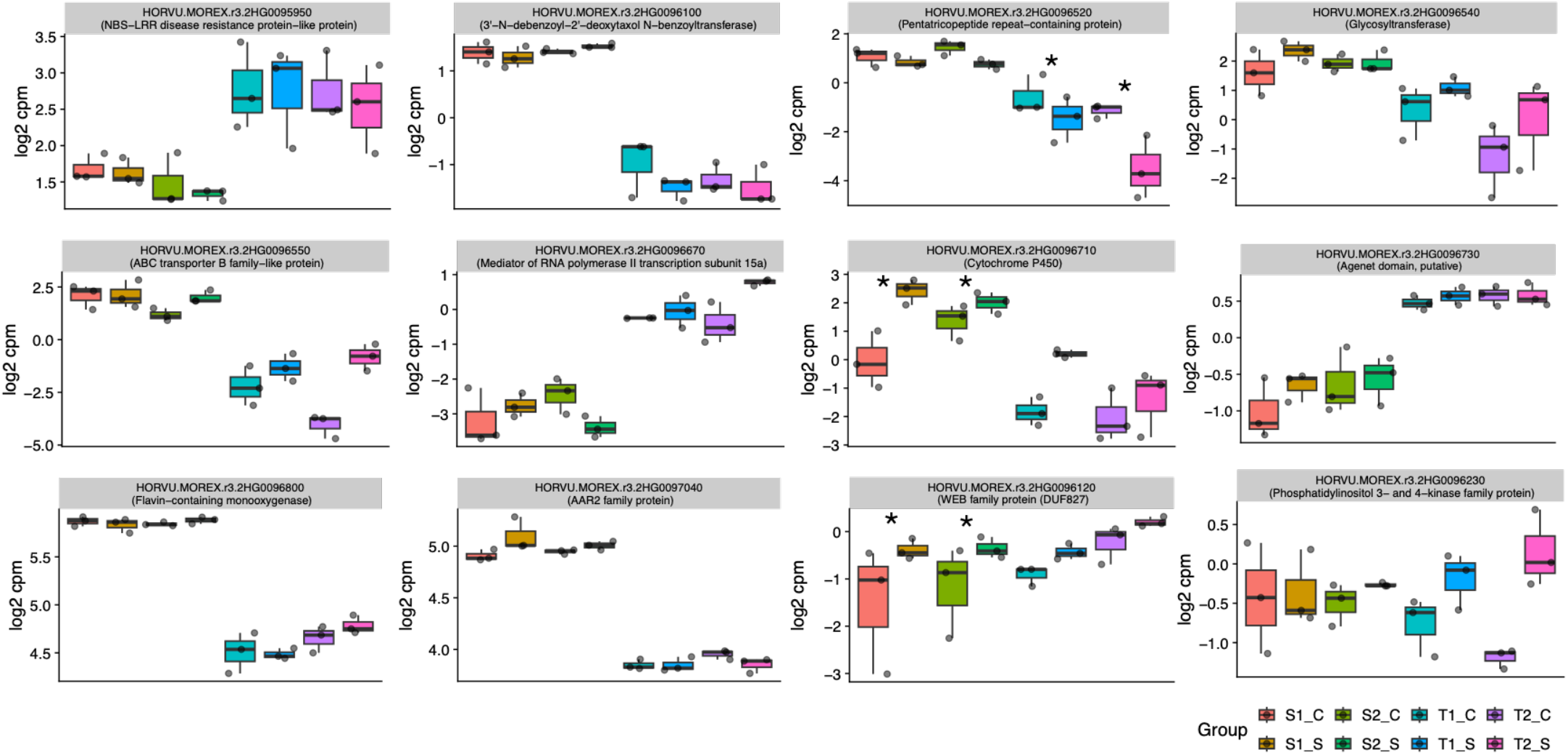
Bulk segregant RNA-Seq shows differentially expressed genes (DEGs) in the QTL interval between 50,000 - 300,000 bp, between the sensitive (S1 and S2) and tolerant (T1 and T2) bulks. Gene expression (Log2 counts per million (CPM) was assessed under control and 100 mM NaCl salt treatment for 3 days. Asterisk indicates a significant difference in the response to salt treatment compared to control.

**Table 4.** List of candidate genes for *QArd.ClSa-2H* and *QSrl.ClSa-2H* between 500,000 – 3,000,000 bp on chromosome 2H.

| Rank | Gene ID | Description | Wheat Gene ID | Rice Gene ID | Arabidopsis Gene ID |
|---|---|---|---|---|---|
| 1 | HORVU.MOREX.r3.2HG0096800 | Flavin-containing monooxygenase (YUCCA 5) | TraesCS2D02G012100.1 | Os04g0128900 (YUCCA7) | AT5G43890 |
| 2 | HORVU.MOREX.r3.2HG0096550 | ABC transporter B family-like protein | TraesCS2B02G012100 | Os05g0548500 (ABC19) |  |
| 3 | HORVU.MOREX.r3.2HG0096770 | Cellulose synthase, putative (DUF1644) | TraesCS2A02G011300.1 | Os01g0612600 | AT3G24740 |
| 4 | HORVU.MOREX.r3.2HG0096540 | Glycosyltransferase | TraesCS2B02G012000 | Os01g0697100 |  |
| 5 | HORVU.MOREX.r3.2HG0096710 | Cytochrome P450 enzyme 71D11 |  |  |  |
| 6 | HORVU.MOREX.r3.2HG0096520 | Pentatricopeptide repeat-containing protein (PPR) |  | Os04t0118700 | AT3G57430 |

## DISCUSSION

Barley is the most salt-tolerant cereal crop compared to other major cereals (Munns and Tester, 2008). While considerable research has focussed on understanding the mechanisms underlying salt tolerance in shoots, the molecular and physiological mechanisms governing root responses remain comparatively poorly understood (Shelden and Munns, 2023). Identifying the molecular mechanisms that contribute to the maintenance of root growth and function during salt stress is important for improving seedling establishment and crop performance in saline soils.

Previous studies showed that Clipper maintains root elongation under salt stress and accumulates higher Na^+^ in the roots than the more sensitive genotype Sahara 3771 (Shelden *et al*., 2020; Shelden *et al*., 2013). These findings suggest that Na^+^ accumulation may contribute to osmotic adjustment and maintenance of water uptake in Clipper, rather than directly inhibiting growth, together with the synthesis of compatible solutes (Shelden *et al*., 2016). Consistent with this hypothesis, salt-tolerant accessions from a global barley core population maintained greater root biomass, which was positively correlated with root Na^+^ content (Tu *et al*., 2021).

### Root responses to salinity and bulk segregant RNA-seq analysis

In this study, we used the parental genotypes, Clipper and Sahara 3771 and144 doubled haploid (DH) lines derived from a Clipper x Sahara 3771 cross to evaluate phenotypic variation in root traits under control and saline conditions. Root traits exhibited quantitative variation in the population, with approximately normal distributions (Figure 1, Figure S1). To minimise background genetic variation in non-targeted traits, we used a pooling strategy to generate tolerant and sensitive bulks for RNA sequencing (Michelmore *et al*., 1991). Bulked segregant RNA-Seq analysis was then used to identify genes differentially expressed between the two groups. This strategy has been successfully used in other studies to identify candidate genes underlying stress adaptation traits, including waterlogging and chilling tolerance in maize (Du *et al*., 2017; Yan *et al*., 2023).

We identified 652 differentially expressed genes in tolerant bulks and 646 in sensitive bulks with 306 overlapping genes, in response to salt stress (Figure 3B). Consistent with previous studies of salinity responses in cereals, salt-responsive genes were enriched for glutathione metabolism, MAPK signalling and phenylpropanoid biosynthesis pathways (Figure 4A and 4B). Glutathione plays a central role in stress tolerance by maintaining redox homeostasis and detoxifying reactive oxygen species (Bose Mazumdar and Chattopadhyay, 2025). MAPK signalling pathways mediate the perception and transduction of stress signals, regulating downstream responses including hormone biosynthesis, transporter activity, and transcriptional reprogramming (Shi *et al*., 2026). Phenylpropanoid-derived compounds, including lignin and suberin, are important components of cell walls and apoplastic barriers that influence water and ion movement. Consistent with this role, previous studies reported enrichment of suberin-related metabolites in Sahara 3771 and lignin metabolites in Clipper (Ho *et al*., 2020), with genotypic differences also reported in wheat (Dissanayake *et al*., 2024). These pathways have established roles in oxidative stress protection and stress signalling, indicating that lines in both bulks activated core salinity-response pathways despite exhibiting contrasting root-growth phenotypes. Flavonoid biosynthesis showed the highest enrichment in tolerant relative to sensitive roots. Flavenoids are polyphenolic secondary metabolites that contribute to antioxidant defence and abiotic stress tolerance in plants, including barley (Mia *et al*., 2026).

In contrast, genes showing a root-length-dependent response to salinity were enriched for transcription factor activity and regulation of RNA biosynthesis, suggesting that differences between tolerant and sensitive lines involve alterations in transcriptional regulatory networks (Figure 4D). This pattern is consistent with the complex genetic architecture of root development and stress adaptation, where changes in regulatory pathways can have substantial downstream effects on root growth and morphology.

### Integration of QTL mapping and BSR-Seq identifies candidate genes

QTL analysis identified loci associated with root morphology, biomass and ion homeostasis, primarily on chromosome 2H but also on chromosomes 1H and 6H. Several of the loci identified in this study overlap genomic regions associated with salinity tolerance in barley. In particular, the chromosome 1H shoot K^+^: Na^+^ ratio co-localises with the *HvNax4/HvCBL4 (HvSOS*3) which controls Na^+^ exclusion under saline conditions (Rivandi *et al*., 2011). Chromosome 2H has also been repeatedly implicated in salinity tolerance across diverse mapping populations, including a QTL for root dry weight identified in the Steptoe x Morex population, although at a different position from the loci detected here (Nguyen *et al*., 2013).

Using a legacy Clipper x Sahara 3771 linkage map, we mapped QTLs for seminal root length and average root diameter between 0 and 10 cM on the short arm of chromosome 2H cM. The Clipper haplotype was associated with longer, thinner roots under both control and saline conditions, whereas the Sahara 3771 haplotype was associated with shorter, thicker roots, especially under salt treatment. To further investigate this region, we designed and applied SNP assays and revised the linkage map of chromosome 2H. This improved the map by replacing existing markers (restriction fragment length polymorphisms and simple sequence repeats) that could not be anchored to the genome assembly and that had missing data. With QTL analysis using the improved map, it became evident that a single QTL (estimated to be at 2.1 cM) affect both root length and diameter (especially under saline conditions), possibly by influencing the balance between longitudinal and radial root growth.

While a similar short-root phenotype has previously been reported in barley under salt stress, the associated QTL was located on chromosome 7H (Xue *et al*., 2017). Identification of a distinct root architecture QTL region on chromosome 2H in the present study therefore reveals an additional genomic region contributing to variation in root architectural responses to salinity in barley.

Improvement of the linkage map also made it possible to anchor the genetic map to the barley genome assembly. The 2.1 cM position corresponded with a 1.3 Mb region (between 1.2 and 2.5 Mb). To allow for the causal gene to be in an adjacent (distal or proximal) region, we conservatively defined a 2.5 Mb region (between 0.5 and 3.0 Mb) as the candidate region. To identify candidate genes within this region, we drew on information from RNA sequence analysis. Of 76 predicted genes in the interval, only a small number exhibited differential expression, sequence polymorphisms with potentially moderate or high impact, or both (Table S10, Table 2-4).

Bulked segregant RNA-seq identified 12 differentially expressed genes within the interval, 10 of which were differentially expressed between long- and short-root phenotypes, and four of which genes were differentially expressed in response to salt treatment in long and/or short roots, respectively (Table 2). Thirty-one genes in the interval were found to contained polymorphic variants with high or moderate impact (Table 3). Six genes were prioritised for further investigation based on expression profiles, sequence variation and predicted biological function. These genes represent strong candidates for regulating root architectural responses to salinity and provide a foundation for future functional validation studies.

### Auxin-associated genes as candidates for root length and diameter

Among the candidates identified within the 2H QTL interval, two genes associated with auxin biosynthesis and transport are particularly compelling given the central role in auxin homeostasis in regulating root elongation and architecture (Korver *et al*., 2018).

The first candidate, HORVU.MOREX.r3.2HG0096800, encodes a flavin-containing monooxygenase of the YUCCA (YUC) family and is homologous to YUCCA genes in rice and Arabidopsis (reference). YUCCA flavin-containing monooxygenases catalyse the conversion of indole-3-pyruvic acid (IPA) to indole 3-acetic acid (IAA), the final and rate-limiting step in the auxin biosynthesis pathway (Vijayanathan *et al*., 2025). The importance of YUCCA-meditated auxin biosynthesis in root development is illustrated in rice where *YUC8/REIN7*, which promotes auxin production and inhibits epidermal cell elongation, results in shorter, swollen roots (Qin *et al*., 2023). Furthermore, increased expression of YUCCA genes under drought and salinity have been associated with enhanced auxin biosynthesis and increased stress tolerance (Marzi *et al*., 2024). Given the central role of auxin in regulating root cell division and elongation, HORVU.MOREX.r3.2HG0096800 represents a strong candidate underlying *QArd.ClSa-2H* and *QSrl.ClSa-2H*.

Expression profiling provides additional support for HORVU.MOREX.r3.2HG0096800 as a candidate gene. Transcript abundance was 2.4-fold lower in lines carrying the Clipper allele compared to those carrying the Sahara 3771 allele (log_2_FC = −1.28, Table 2, Figure 7). In addition, the gene contained a missense polymorphism resulting in an Asp (D) to Asn (N) substitution at amino acid position 65. The Asp residue is highly conserved among homologous proteins in barley, rice and Arabidopsis and in the Clipper allele, suggesting that the variant warrants further investigation for effects on protein function. Notably, the Sahara 3771 allele is associated with the shorter, thicker root phenotype, raising the possibility that increased expression of this auxin biosynthesis gene alters local auxin homeostasis in the root apex and consequently influences the balance between longitudinal and radial root growth.

Although expression of HORVU.MOREX.r3.2HG0096800 was not significantly altered by salt treatment, the strong difference between allelic classes suggests that constitutive differences in auxin regulation, rather than salt-induced transcription of this gene, may contribute to the contrasting root phenotypes. This distinction is important because root responses to salinity involve not only changes in auxin biosynthesis but also changes in auxin redistribution within the root. Salt-induced inhibited of root elongation has been associated with reduced auxin accumulation (Liu *et al*., 2015) which appears to result from altered polar auxin transport, as salinity affects the localisation of auxin transporters within the root apex (Jiang *et al*., 2016).

The second candidate gene, HORVU.MOREX.r3.2HG0096550, is predicted to encode an ATP-binding cassette subfamily B (ABCB) transporter. Members of the ABCB transporter family are key components of the polar transport of auxin and have important roles in the regulation of root growth, lateral root development and root architecture (Devi *et al*., 2025; Geisler *et al*., 2017). They have also recently been shown to transport other phytohormones including brassinosteroid (Ying *et al*., 2024). ABCB transporters also function cooperatively with PIN transporters to regulate the spatial distribution of auxin in root tissues (Mellor *et al*., 2022). In barley, the ABCB transporter HvABCB25, contributes to aluminium tolerance, and its overexpression alleviates aluminium-induced inhibition of root elongation (Liu *et al*., 2021), demonstrating that ABCB transporters can influence root growth responses to abiotic stress in cereal crops.

In the present study, HORVU.MOREX.r3.2HG0096550 was more highly expressed in root tips of lines carrying the Sahara 3771 allele than those carrying the Clipper allele (Figure 7). Expression increased under salt treatment in lines carrying the Clipper allele. As observed for the YUCCA-like candidate, the clearest expression difference was associated with genotype rather than salt-treatment. If HORVU.MOREX.r3.2HG0096550 functions in auxin transport, variation in its expression or activity could alter auxin distribution within the root apex and contribute to differences in root elongation and radial growth. Consequently, this gene also represents a plausible candidate underlying chromosome 2HS QTL interval.

Together, these results suggest that variation in auxin homeostasis may contribute to the contrasting root architectures of Clipper and Sahara 3771. Both the YUCCA like and the ABCB transporter were more highly expressed in the root tips of lines carrying the Sahara 3771 allele which is associated with shorter, thicker roots. As these genes occupy complementary positions with the auxin pathway, affecting auxin biosynthesis and transport respectively, coordinated variation in their activity could influence auxin abundance and distribution within the root apex, altering the balance between longitudinal and radial root growth. However, because the QTL effects were detected under both control and saline conditions, these loci may regulate constitutive differences in root development rather than a salt-specific response. Functional validation of both, together with measurements of auxin distribution and root cellular anatomy, will be required to test this hypothesis.

### Other gene candidates

Although the two auxin-associated genes provide biologically compelling hypotheses for the chromosome 2H root phenotype, several other genes within the QTL interval warrant consideration. HORVU.MOREX.r3.2HG0096540 encodes a UDP-glucosyltransferase (UGT), that may play a role in the glycosylation of metabolites, including phytohormones, as has been shown in other species (Wang *et al*., 2023). Through their effects on hormone homeostasis, UGTs have been implicated in plant growth, development and responses to abiotic stress. Consequently, variation in the expression or activity of this gene could influence root development either directly or through modification of hormone signalling pathways.

Another candidate, HORVU.MOREX.r3.2HG0096770, is predicted to encode a DUF1644-containing protein associated with cellulose synthase. Cellulose synthases are essential for cell wall formation and root cell expansion (Polko and Kieber, 2019). Consistent with this, RNAi-mediated silencing of a cellulose synthase gene in barley resulted in slower root growth, linked with a shorter elongation zone and a significant reduction in root system size (Lou *et al*., 2022). Variation in the expression or function of a cellulose synthase gene could therefore contribute to differences in root elongation.

A further notable candidate, HORVU.MOREX.r3.2HG0096710, which exhibited large difference in expression between the contrasting root phenotypes, is predicted to encode a member of the cytochrome P450 superfamily. Members of the P450 superfamily have previously been associated with primary and lateral root development (Julkowska *et al*., 2017; Tu *et al*., 2021). Several cytochrome P450 families participate in hormone metabolism. For example, CYP79B2 and CYP79B3 convert tryptophan to indole-3-acetaldoxime (IOX), linking P450 activity with pathways contributing to auxin metabolism (Hull *et al*., 2000), while other P450 enzymes regulate ABA homeostasis through ABA catabolism (Kushiro *et al*., 2004)). Furthermore, disruption of CYP83B (*Sur2*) in Arabidopsis increases free IAA levels and promotes lateral root development (Barlier *et al*., 2000). Although these observations make HORVU.MOREX.r3.2HG0096710 an interesting candidate, its biochemical function cannot be inferred from its general P450 annotation, and identification of its CYP subfamily will be required to establish a role in hormone metabolism.

HORVU.MOREX.r3.2HG0096520 is predicted to encode a pentatricopeptide repeat protein (PPR). These proteins are typically targeted to the mitochondria or chloroplast, where they regulate RNA processing, editing and translation and have been implicated in abiotic stress tolerance. Notably, a PPR gene was identified within a QTL associated with root growth under salinity stress in barley (Cope *et al*., 2022), providing support for a possible role in the regulation of root responses to salinity.

Together, these candidate genes identified within the QTL interval represent several plausible biological mechanisms that could influence root architecture, including auxin homeostasis, hormone modification, and cell wall biosynthesis. The development of thicker roots under salinity may represent an adaptive response that alters radial transport pathways, ion uptake, or root hydraulic properties. Further functional characterisation of these candidate genes will be required to identify the causal gene and establish whether auxin-mediated changes in root architecture contribute directly to improved tolerance of saline conditions.

## Conclusion

In summary, the integration of QTL mapping, BSR-sequencing and sequence variant analysis identified a small number of candidate genes in a region of barley chromosome 2H that affects root architecture traits. Several candidates have functions consistent with roles in root development and stress responses, including genes involved in auxin biosynthesis and transport. These genes represent plausible candidates for the observed differences in longitudinal and radial root growth between Clipper and Sahara 3771, potentially by affecting auxin homeostasis. These findings provide new insight into the genetic regulation of root architecture and establish a foundation for future studies investigating the role of root traits in adaptation to saline soil.

## ACKNOWLEDGEMENTS

We would like to acknowledge the contribution of the South Australian Genomics Centre (SAGC) in the generation of data used in this publication. The SAGC is supported by its founding partner institutes and by BioPlatforms Australia as part of the Australian Government’s National Collaborative Research Infrastructure Strategy (NCRIS). We also acknowledge the Australian Centre for Plant Functional Genomics for providing the mapping population and associated data. The authors wish to thank to Professor Matthew Tucker and Dr Nick Collins for helpful discussions and Dr. Adriane Piechatzek, Dr Jessey George and Ms. Jodie Kretschmer for technical assistance. MS acknowledges financial support from the Australian Research Council (DE140100575) and Adelaide University Mortlock Bequest.

